# Assessment of Leaf-Litter Invertebrate Biodiversity Using High Throughput Sequencing

**DOI:** 10.64898/2026.04.09.717398

**Authors:** Anibal H. Castillo, Shoshanah Jacobs, Dirk Steinke, M. Alex Smith

**Author notes:** **Corresponding Author**: Anibal H. Castillo, **Email address**.

## Abstract

Leaf litter ecosystems and their fauna are largely understudied, despite their critical ecological roles. Here, we investigate challenges associated with estimating biodiversity in terrestrial leaf litter. Current methodologies for biodiversity assessment are fraught with limitations, amongst the most significant is a decline in taxonomic expertise, complicating the process of species identification and the significant costs associated with species-level morphological identifications. DNA barcoding employs the mitochondrial gene cytochrome *c* oxidase I (COI) to identify animal species, and DNA metabarcoding facilitates the identification of multiple species without necessitating taxonomic expertise. Recent studies indicate that environmental DNA (eDNA) may exhibit greater sensitivity compared to traditional methods. To test whether these methods work in a real-world application, we sampled leaf litter across a temperate forest/field ecotone. Leaf litter was dried, ground and processed to extract environmental DNA. We evaluated the DNA extraction protocols to test their relative efficacy. We found that the Qiagen Blood and Tissue Kit was the most effective at recovering invertebrate diversity and that there were notable differences in biodiversity between forest and field habitats. Temperature emerged as a significant factor influencing the composition of the communities observed. Our methodology is applicable across various environments for efficient biodiversity assessment and might be particularly beneficial for monitoring pests and invasive species. Our approach offers a cost-effective and timely alternative to conventional biodiversity assessment methods and underscores the significance of accurate assessment methodologies for leaf litter communities.

## Introduction

Estimating biodiversity is as challenging as it is important, especially during the Anthropocene [1,2]. A good example of a challenging and largely understudied ecosystem is found in the terrestrial leaf litter. While leaf litter communities are vital for ecosystem services and other functions [3–6], the species that comprise them are largely unknown, mainly because approaches are inconsistent, and very few methods exist to attempt more comprehensive surveys.

Evaluating the animal biodiversity found in the leaf litter includes collection, storage, faunal extraction, sorting and identification [7]. While each one of these stages has limitations and biases, collecting and storing leaf litter to target invertebrate biodiversity is relatively simple. For example, in one widely accepted standardized protocol, a predetermined small section, typically 25 cm x 25 cm, is collected wearing gloves and then kept cold or frozen pending further processing [8]. However, once collected, many factors influence the subsequent estimation of biodiversity, and no single sampling method can comprehensively assess all taxa encompassing the leaf litter’s terrestrial biodiversity [9]. Montgomery et al., (2021) recently reviewed seven collection methods that, in their taxonomic overlap, might offer a better assessment of terrestrial biodiversity. However, the challenge of achieving a comprehensive estimate with a single approach remains largely untenable. An assessment of a sample’s biodiversity is only as good as one’s ability to identify the taxa collected. An evaluation of an area’s diversity, where specimens can only be keyed to order, will not be as good as one where we can identify them to genus, let alone species (although see [11]. Thus, the extent of available taxonomic expertise in each collected group limits the ultimate evaluation. Especially for soil invertebrates common in the leaf litter (such as nematodes, earthworms, and mites), taxonomic expertise is dwindling, and for other taxa like proturans, diplura and symphylans, it is already scarce [12]. This is exacerbated by small adult size, the prevalence of cryptic species, which can result in underestimates of morphologically-derived species richness, and changes in morphology between sexes or across life-history stages. Molecular methods of species identification, including DNA barcoding and metabarcoding, pose a potential solution to these problems and may represent a faster and more cost-effective way to assess terrestrial biodiversity, helping to mitigate its decline and manage its recovery.

DNA barcoding uses a standardized fragment of the mitochondrial gene cytochrome *c* oxidase I (COI), to detect the presence of animal species in various environments [13]. Over the past twenty years, it has fueled a better understanding of community networks [14–16]. Its extension, DNA metabarcoding, provides a “genetic profile” that can be screened for multiple species of interest without requiring taxonomic expertise. This approach, which utilizes High-Throughput Sequencing (HTS) technology, can also be applied to soil arthropod profiling (Oliverio et al., 2018). In fact, several studies (Allen et al. (2021; Weber et al. (2024) showed that environmental DNA (eDNA) as a source for metabarcoding can be more sensitive than conventional methods when detecting invertebrate species (see also Robinson et al., 2022), opening up new sources of samples, particularly from leaf washes and drying and grinding of leaves. The critical stage of linking the produced sequences to formal names is dependent on the existence of a proper reference library [20].

### Opportunity

To date, when leaf-litter diversity has been analyzed via HTS – it has been via traditional collection methods (like Winkler sifting, e.g., Yang et al., 2014) where the arthropods are first extracted from the matrix and then the HTS protocol is applied to the emergent arthropods (i.e., after the rate-limiting sorting steps outlined earlier). However, leaf-litter animal biodiversity has yet to be directly explored using HTS when these rate-limiting steps are removed. In an intriguing parallel, Krehenwinkel et al. (2022) extracted arthropod DNA from store-bought tea using a CTAB protocol. While it may seem contradictory that they chose a protocol often used to extract plant DNA [23] the authors designed primers to minimize the amplification of plant DNA, specifically to target arthropod DNA within and upon the collected plant material. In addition, environmental assessment methods using DNA have not been extensively employed to monitor plants [24,25] because they may not always release detectable amounts of DNA [26]. The novel approach pioneered by Krehenwinkel et al allowed the recovery of ecologically and taxonomically diverse arthropod communities from the tea, many specific to their host plant and its geographical origin (Krehenwinkel, Weber, Künzel, & Kennedy, 2022). The same research group (Krehenwinkel, Weber, Broekmann, Melcher, et al., 2022) applied a similar approach to herbarium leaves, enabling them to study temporal changes in arthropod communities. Atypically for eDNA, arthropod DNA from dried plants showed remarkably high temporal stability, opening the possibility for studies on the ecology of invertebrates using the plant material they inhabit, such as leaf litter. More recently, the same research group studied arthropod biodiversity on plant leaves through freeze-drying and grinding prior to HTS [16]. Here, we expand their approach to determine if HTS is feasible for studying invertebrate communities in a leaf-litter environment. We have designed an approach that involves drying and grinding leaf litter into a fine powder from which we extract environmental DNA, used to identify invertebrate taxa using conventional metabarcoding pipelines. As a proof of concept, we tested this new approach with three different extraction kits and then directly assessed the terrestrial biodiversity across a forest-to-field ecotone. Using this novel approach, we wanted to ask the questions: how does extraction protocol affect the recovery of the arthropod community, and can we use the best protocol to analyze how arthropod leaf-litter diversity varies across a temperate forest/field ecotone?

## Materials and Methods

### Experimental Design

We sampled leaf litter in both a small woodlot and an adjacent early-succession field on the University of Guelph campus (Dairy Bush, Guelph, Ontario, Canada) in late October 2022, following the SoilBON protocol [8]. We performed half of the sampling in the field site (top-left white cross in **Error! Reference source not found.**) and the other half in the woodlot (bottom-right cross in **Error! Reference source not found.**). Significantly different temperature regimes characterize both areas during the growing season (Smith 2024). We collected five samples in each area using a 25 cm by 25 cm quadrat, where the design follows an axis that aligns with the temperature gradient (Smith 2024) (**Error! Reference source not found.**). This step included all plant material (e.g., grass and plants) to ensure all eDNA could be recovered in the leaf-litter powder. Temperature measurements were taken every 10 minutes at ground level throughout the year using Onset HOBO MX2201 Pendant Wireless Temperature Data Loggers (**Error! Reference source not found.**) [28].

### Drying and Grinding of the Leaf Litter

We did not wash or alter the leaf litter before processing (Krehenwinkel, Weber, Künzel, & Kennedy, 2022). We dried the leaf litter at 56 °C and weighed it regularly until the weight stopped dropping (24 hours). Then, we ground the samples into a fine powder using an IKA Tube Mill control (IKA, Breisgau, Germany) at 25,000 rpm for three minutes twice [29]. Additionally, we combined an aliquot of the five samples from each site to form a sixth sample. We compared sequencing results from this mixture to those of the five original samples to determine if it was representative of the environment. If so, the cost and effort of sequencing could be reduced by 80%.

### Processing Costs

The Biological Survey of Canada (1994) estimated that processing a terrestrial arthropod inventory at any individual site requires approximately 504 hours each month when utilizing a standard sampling protocol. If biology undergraduate students are employed at an updated rate of US$10.95 per hour, the monthly expense for processing the collected materials at the site would total US$5,518.65. Thus, for a single-site inventory in most regions of Canada, processing from April to October over a duration of seven months would result in a total expenditure of US$38,630.56. In comparison to these processing costs, other expenses such as sampling supplies and travel are relatively minor. It is essential to recognize that processing costs are incurred only until species identification becomes feasible [30].

After this point, expenses related to the already limited taxon-specific expertise increase and are challenging to evaluate. Regarding laboratory expenses, [31] projected that costs associated with metabarcoding range from US$240 to US$415 per sample, based on the now-defunct Roche GS FLX ‘454’ sequencer. However, the Illumina sequencing technology used in our research is significantly less expensive, approximately 54.50 USD [32] (**Error! Reference source not found.**). In terms of processing efforts Ji et al. (2013) assessed that the “active workload” (comprising lab and computer work) related to metabarcoding necessitates approximately one-quarter of the effort needed for visual sorting of indicator taxa into morphospecies among arthropod taxa. For the microscopic taxa commonly found in ground-level samples, metabarcoding provides a notably greater advantage regarding time. Furthermore, while costs associated with metabarcoding increase on a per-sample basis, standard biodiversity expenses escalate on a per-specimen basis. This limitation elucidates why traditional biodiversity censuses focus solely on indicator taxa [31]. Consequently, a typical molecular laboratory, staffed by a small team, can process hundreds of whole samples annually from anywhere in the world, achieving a data production rate and breadth that proves unattainable with traditional methodologies. Still, metabarcoding approaches must be conducted in synergy with traditional taxonomy work, as the former is heavily reliant on a robust reference library that can only be provided by the latter.

### DNA Extraction, PCR Amplification, and Primer Choice

We extracted total DNA from leaf-litter powder using three different commercial kits on each sample: a DNeasy® PowerSoil® Pro Kit (Qiagen, Hilden, Germany), a DNeasy® Blood and Tissue Kit (Qiagen, Hilden, Germany), and a Quick-DNA^TM^ Plant/Seed Miniprep Kit (Zymo Research, Irvine, CA, USA) (**Error! Reference source not found.**). The same amount of tissue was not added to each kit, which were used according to the manufacturers’ instructions.

We measured the quality and yield of the DNA extractions using a Qubit^TM^ dsDNA HS Assay Kit (Invitrogen, Life Technologies, Carlsbad, CA, USA) and on a 1% agarose gel. We performed PCR amplifications using a two-step fusion primer PCR protocol [33]. We amplified a 421 bp region of the cytochrome *c* oxidase subunit I (COI) during the first PCR step using the BF3 + BR2 primer set [34]. We performed PCR reactions in a 25 μL reaction volume, with 0.5 μL DNA, 0.2 μM of each primer, and 12.5 μL PCR Multiplex Plus buffer (Qiagen, Hilden, Germany). We used a Veriti thermocycler (Thermo Fisher Scientific, MA, USA) under the following cycling conditions: initial denaturation at 95 °C for five min; 25 cycles of 30 sec at 95 °C, 30 sec at 50 °C and 50 sec at 72 °C; and a final extension of five min at 72 °C. PCR success was checked on a 1% agarose gel. We used one μL of PCR product as a template for the second PCR, where we added Illumina sequencing adapters using individually tagged fusion primers [33]. We used the same thermocycler conditions as in the first PCR but increased the reaction volume to 35 μL by adding water, reduced the cycle number to 20, and increased the extension time to 2 minutes per cycle. We checked PCR success again on a 1% agarose gel. We purified and normalized PCR products using SequalPrep Normalization Plates (Thermo Fisher Scientific, MA, USA, Harris et al. 2010) according to manufacturer protocols. We pooled ten μL of each normalized sample and cleaned the final library using left-sided size selection with 0.76x SPRIselect (Beckman Coulter, CA, USA). This method removes DNA fragments that are smaller than the targeted size, allowing for the selection of larger fragments. At 0.76x, SPRIselect targets fragments longer than 400 bp. The PCR fragment with the primers used is 421 bp. Sequencing was performed at the Advanced Analysis Centre at the University of Guelph using the 600-cycle Illumina MiSeq Reagent Kit v3 and 5% PhiX spike-in.

### Bioinformatic Analyses and Taxonomic Classification

We performed Bioinformatic analysis as outlined in Buchner et al., (2022): after demultiplexing, we used the APSCALE pipeline [35], and taxonomy was assigned to Operational Taxonomic Units (OTUs) using a curated Canadian Reference Library (Pentinsaari et al., in prep – available on BOLD).

### Statistical Analyses

All statistical analyses were executed in R (R version 4.3.3 (2024-02-29), Team, 2020) using RStudio (Version 2025.05.0+496). We conducted sample-based rarefaction analyses to confirm that our sequencing depth was adequate. To visualize the differences in community structure and composition among sites [37], we employed non-metric multidimensional scaling (nMDS) ordination with k=2, utilizing the ‘vegan’ package [38]. The nMDS plot is based on Bray-Curtis distances, using species presence/absence data. All the necessary code and datasets for reproducing these results, including data visualization, are available online at [insert link].

## Results

A total of 20,361,353 paired-end raw COI reads were sequenced (an average of 535,825 reads per sample). After merging, removal of the primer sequences, length selection, quality filtering, and denoising, 16,289,082 reads were used for dereplication and subsequent species identification. Dereplicated raw reads are available at the NCBI Short Read Archive (SRA) under the accession number XX.

All three DNA extraction protocols successfully recovered DNA from the leaf-litter powder (**Error! Reference source not found.**). We found different levels of success in recovering the leaf-litter invertebrate communities according to the protocol used (**Error! Reference source not found.**). Because all extractions came from the same sample, we considered the most successful protocol to be the one exhibiting the highest alpha diversity (number of different taxa).

There was a significant interaction between habitat and DNA extraction method (i.e., not all methods performed equally in all habitats). PowerSoil Pro and Blood and Tissue methods exhibited greater diversity in the field than in the forest, whereas Zymo showed no difference between the habitats. The Blood and Tissue Kit appeared to capture a greater diversity compared to PowerSoil Pro and Zymo (**Error! Reference source not found.**) (p-value 0.023). When analyzed according to different habitats, PowerSoil Pro performed well in the field but less effectively in the forest. Blood and Tissue performed exceptionally well in the field and reasonably well in the forest. In contrast, Zymo performed average in both environments (see **Error! Reference source not found.**). For all three protocols, the field samples resulted in higher DNA yields than the forest samples (***Error! Reference source not found.***), likely because the field samples were collected while the plants were alive. In contrast, the forest samples consisted primarily of dead leaves. An nMDS plot illustrating the beta diversity by extraction type revealed that the Blood and Tissue Kit captured a greater breadth of diversity (**Error! Reference source not found.**).

We anticipated that warmer areas in the field would exhibit greater alpha diversity, and that field and forest communities would be different. An nMDS plot displaying beta diversity by habitat, with vectors corresponding to average air temperature, average maximum temperature, average minimum temperature, soil conductivity, water volume, and soil temperature (**Error! Reference source not found.**), revealed that temperature was the most significant factor explaining the majority of the variance (p = 0.001). Interestingly, an nMDS plot of the samples that includes the taxa illustrated the retrieval of different taxa within three of the most abundant classes we observed (e.g., Arachnida, Insecta and Collembola) across the field and forest environments (see **Error! Reference source not found.**).

## Discussion

To our knowledge, ours is the first study to apply metabarcoding of eDNA to leaf-litter biodiversity assessment. Based on earlier work [16,39–41], we assumed that drying and grinding leaf litter with subsequent DNA extraction would be an effective way to evaluate the resident terrestrial arthropod communities. Here, we successfully applied our approach to distinguish between markedly different invertebrate communities in adjacent field and forest environments.

### Protocols

To find the most optimal method for extracting DNA from leaf litter, we tested three commercial DNA extraction protocols. Based on our results, the Qiagen Blood and Tissue Kit was most successful, as it returned the highest quantities of DNA and recovered the greatest alpha diversity (measured as BINs recovered). This kit represents the most versatile of all three options, as Qiagen PowerSoil Pro is designed for microbial DNA extraction, while Zymo is more specialized for plant DNA extractions.

We predicted that combining replicates before processing would retrieve similar alpha diversity and thereby reduce labor and costs. However, combining aliquots from each replicate did not result in taxonomic coverage representative of the sum of the replicates. This confirms earlier findings that technical replicates are required for eDNA metabarcoding to accurately recover alpha diversity [42].

### Environmental Comparison

Our approach has successfully differentiated between invertebrate communities found in adjacent sites across a forest/field ecotone. Notably, we found that none of the species were shared across the ecotone for the three most common groups we sequenced (Arachnida, Insecta and Collembola, **Error! Reference source not found.**). The section of the field from which we collected samples has not been cut since 2018, and as such, it has been left to develop into a forest through a process of secondary succession. Currently, there are many young tree saplings from numerous species common in the adjacent forest that are emerging and (at writing) are upwards of meters tall (Smith 2024). What will happen to the distinct community of arthropod species we found in the field if it becomes a forest? Will they be replaced by the nearby forest species, become locally extinct, or move to another location? Alternatively, what will happen if the University landowners decide to cut the emergent forest? Our findings in this small methodological case study not only underscore the need for further research but also reflect how subtle abiotic gradients can produce significant biological effects – even in a developed urban ecosystem and hopefully inspire future studies in invertebrate community ecology.

### Applications

Our approach complements comprehensive environmental assessments and can be applied to various types of plant residue. This could include applications such as screening crop samples for pests or invasive species in shipping materials [43–45]. Leaf litter assessments can be conducted in a variety of environments, including grasslands, forests, and agricultural and horticultural fields. However, this approach can only be used when leaves, either as leaf litter or still on the live plant, are available. Additionally, it is conceivable that if the leaf litter has recently fallen, its inhabiting fauna might not have crawled over it and would not have left behind recoverable DNA. The only DNA available would be that of organisms that left it on the leaves while they were part of the canopy.

Compared to other methods of monitoring arthropods in the applied agricultural world, our method is efficient. Residual plant material can be collected quickly, and the entire process takes only a few weeks, primarily consisting of waiting for sequencing results, which can be expedited to a few days. Furthermore, it is cost-effective, as sampling efforts and laboratory work can be less expensive than those required by traditional methods. Additionally, it offers an out-of-season opportunity for sampling, as leaf litter collected in non-active seasons could provide insight into the biodiversity present in the environment during the previous active season.

### Conclusion

We have demonstrated the feasibility of capturing invertebrate communities in leaf litter by drying and grinding only the leaves. This illustrates that the HTS-derived data can be used to assess interactions between these communities. Such a method will be extremely useful in theoretical and applied conditions.

## Acknowledgments

The Natural Sciences and Engineering Research Council and Food from Thought funded the project. We also thank anonymous reviewers for their help in improving earlier versions of this manuscript.

## Figures

**Figure 1.**
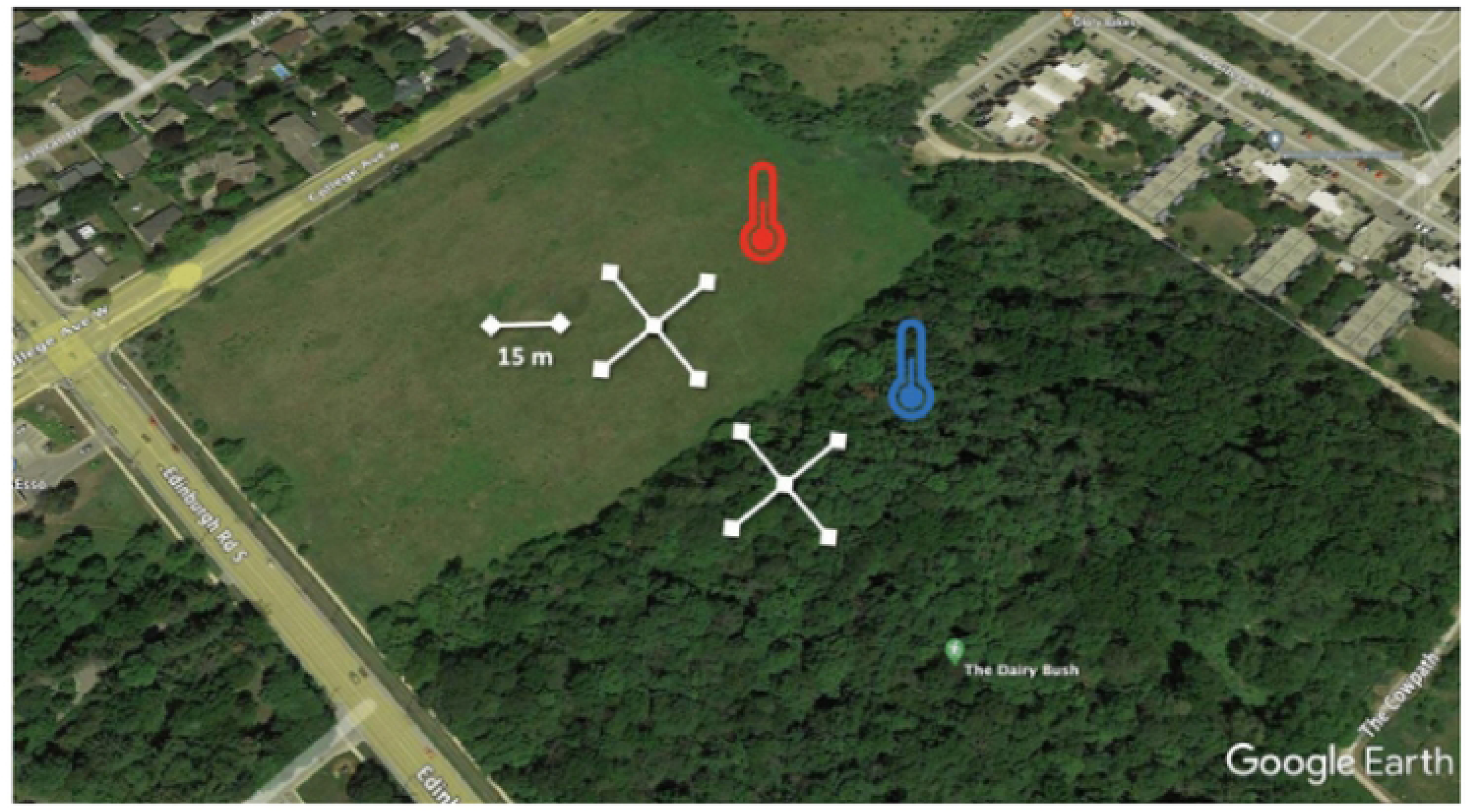
Sampling and experimental design. We sampled leaf litter in the Dairy Bush using the SoilBON protocol [8]. We collected five samples in the field and five in the forest using a 25 cn1 by 25 cm quadrat and spacing each point 15 1n apa1t. The sa1npling points follow an axis along the forest/field ecotone, which has a predictable gradient of temperature and 1noisture.

**Figure 2.**
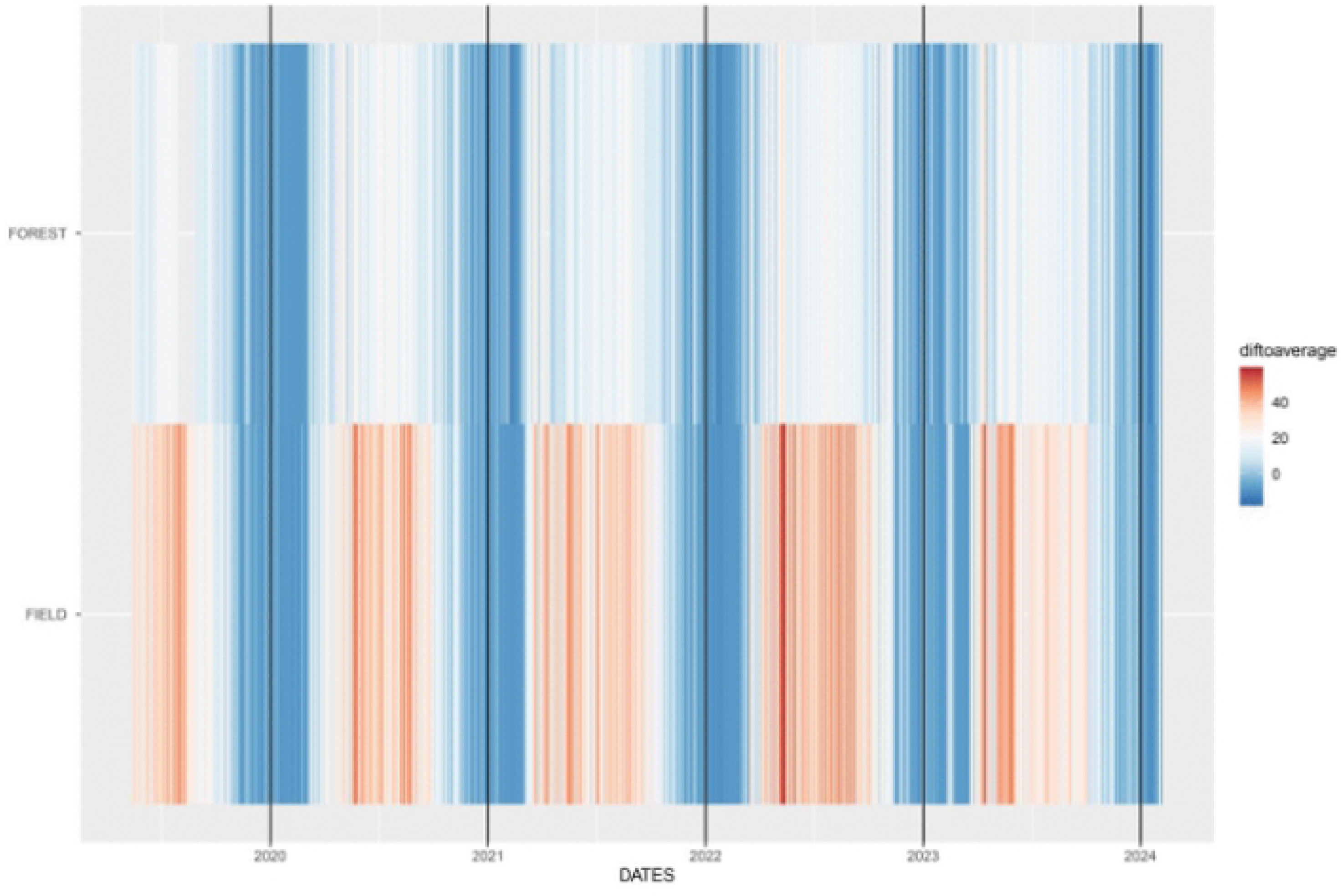
The daily maximum temperature in the Dairy Bush Forest and Field between May 2019 and February 2024, when compared to the annual average temperature in Guelph, Ontario, Canada (7.8 °C). In this plot, each vertical line is a day, coloured by the difference between the average Guelph temperature. The growing season difference between the two habitats and their similarity in the winter are evident.

**Figure 3.**
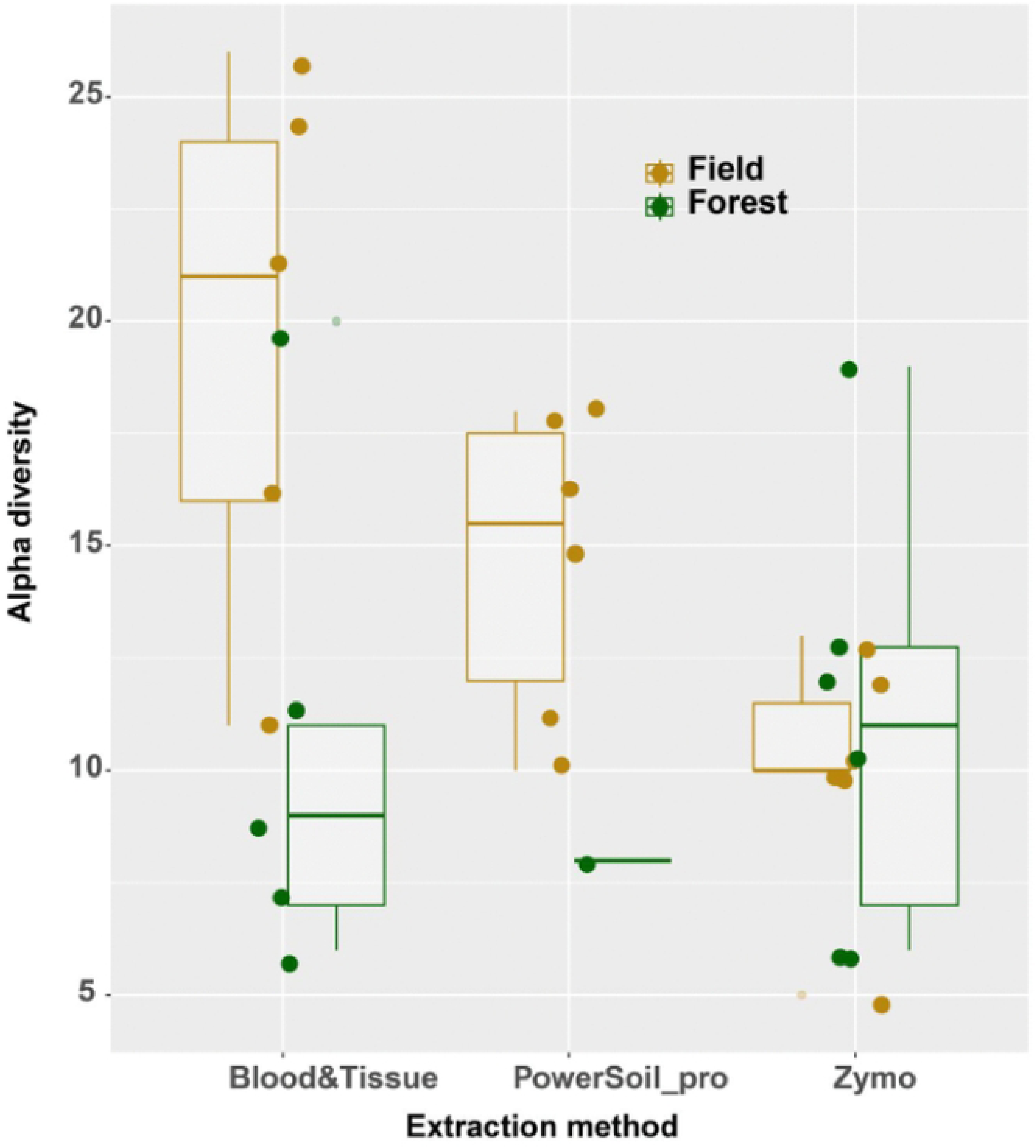
The Blood and Tissue kit captures more diversity than the other two kits tested (p-value 0.023). By habitat, the results indicate that PowerSoil performs well in the field but lags behind in the forest. Conversely, Blood and Tissue performs exceptionally well in the field and reasonably well in the forest. In contrast, Zymo performs average in both environments. Overall, Blood and Tissue is the most effective at reliably recovering the diversity from both environments.

**Figure 4.**
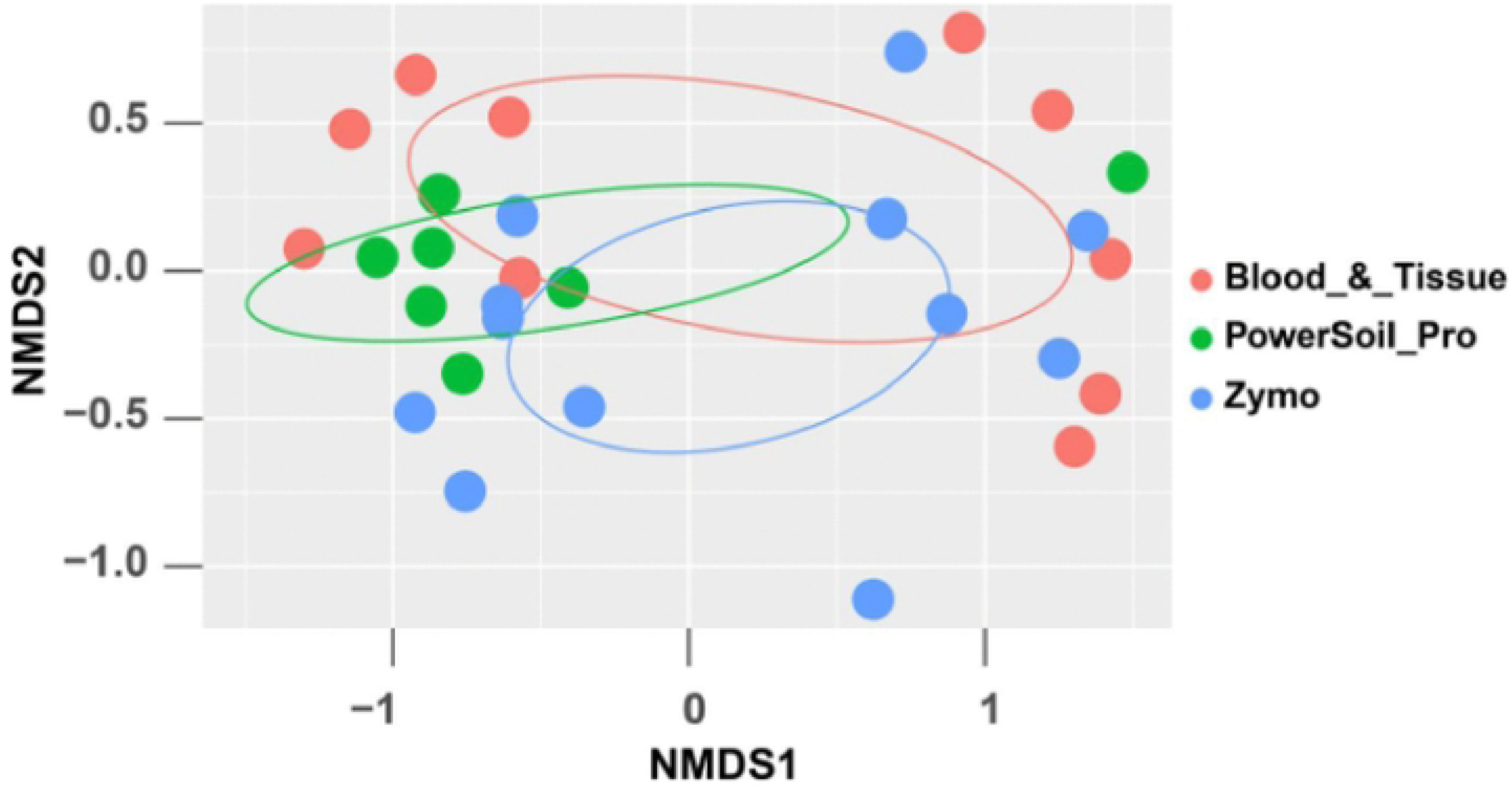
NMDS of beta diversity by extraction type: DNeasy® Blood and Tissue Kit (Qiagen, Hilden, Gem1any); DNeasy® PowerSoil® Pro Kit (Qiagen, Hilden, Germany); and Quick-DNA TM Plant/Seed Miniprep Kit (Zymo Research, Irvine, CA, USA). (stress value: 0.118).

**Figure 5.**
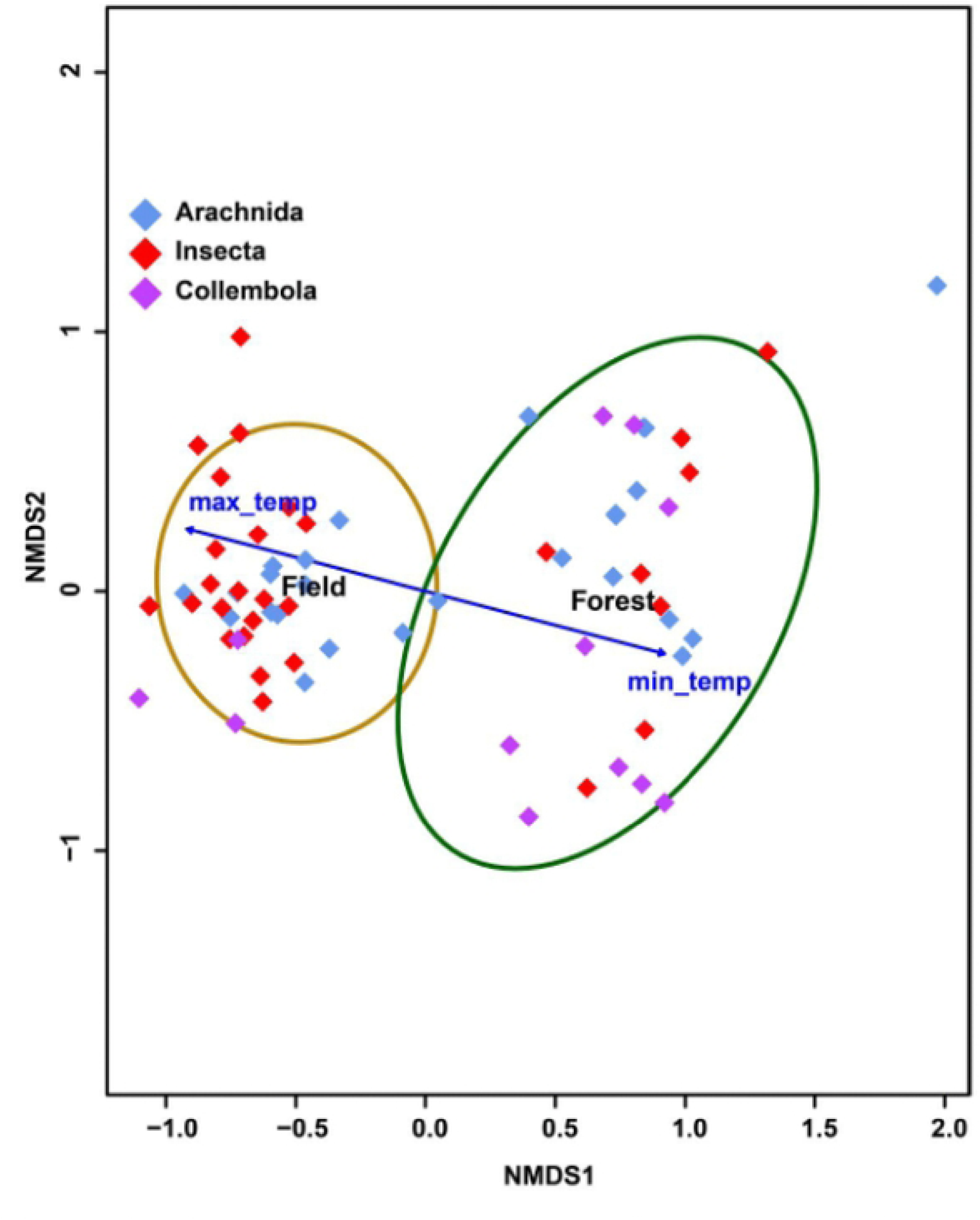
Visualizing community structure across environment types associated with taxa at the Dairy Bush (Guelph, Ontario, Canada) using non-metric multidimensional scaling (nMDS) based on ten samples, five from each of two adjacent. locations. The vectors correspond to average air temperature (avg_ai11en1p), average maximum temperature ( avg_maxtemp), average minimum temperature ( avg_mintemp), soi I conductivity (SC), water volu1ne (:NV), and soil temperature (SoilTemp). (a) The first NMDS di1nension is largely an expression of the abiotic te1nperature and 1noisture environ1nent across the ecotone. The co1n1nunities in both environ1nents (coloured 95% ellipsoids)-types are significantly different (ANOSIM statistic R: 0.877; significance: 0.00 I; stress value: 0.1 17).

**Table 1.**
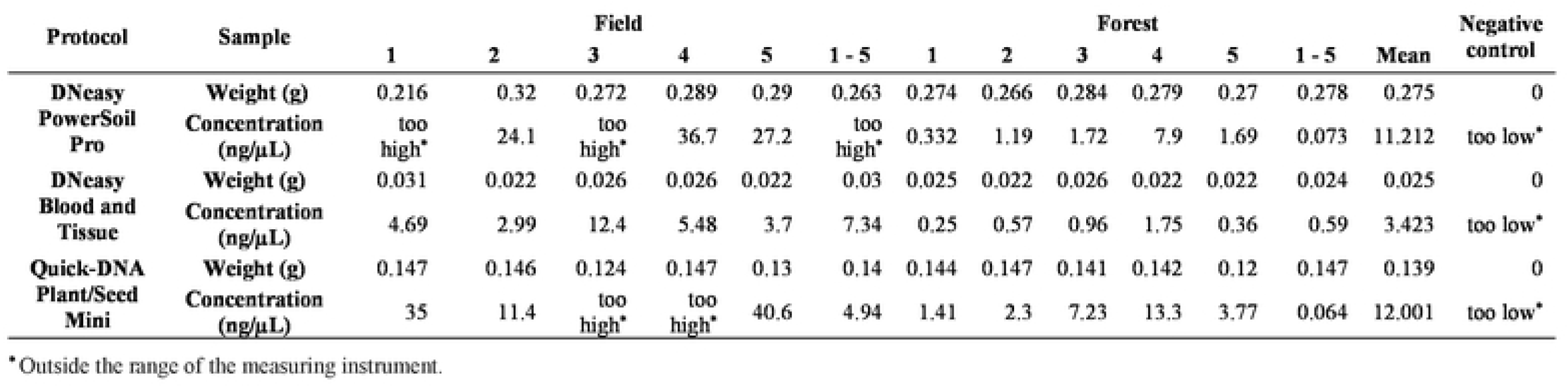
Total DNA extraction yield for each of the three extraction protocols. (ps) DNeasy PowerSoil Pro; (bt) DNeasy Blood and Tissue; (zm) Quick-DNA TM Plant/Seed Mini.

**Table 2.**
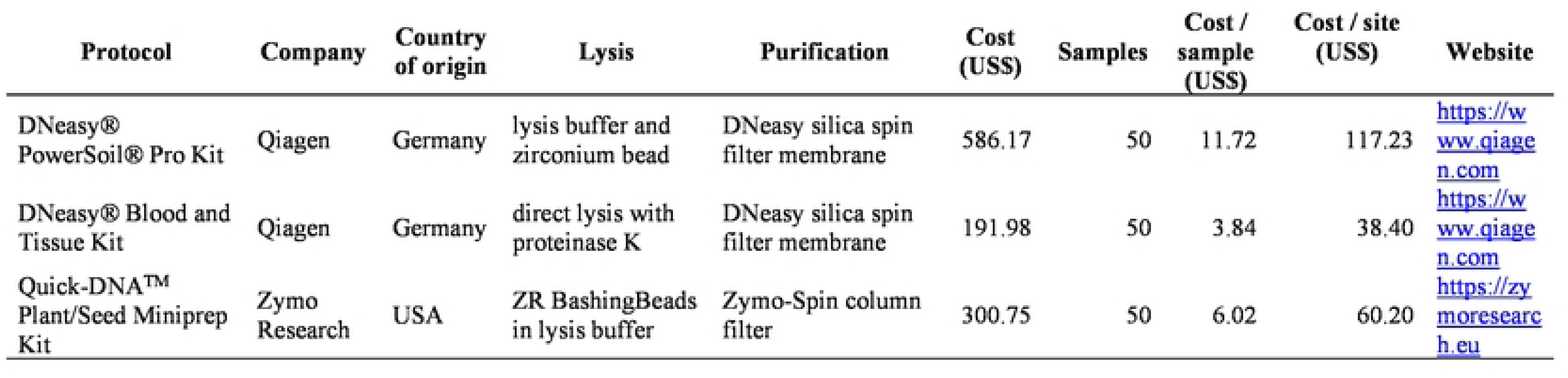
Processing costs for the three extraction protocols: DNeasy® PowerSoil® Pro Kit, DNeasy® Blood and Tissue Kit, and Quick-ONATM Plant/Seed Mini prep Kit, used in this study.

